# A transient feature of the inferior olive supports the development of cerebellar internal models

**DOI:** 10.64898/2025.12.06.692744

**Authors:** Angela M. Richardson, Greta Sokoloff, Mark S. Blumberg

## Abstract

The inferior olive (IO) supports motor learning by supplying the cerebellum with critical sensory and motor input. In adult rats, that input includes externally generated limb stimulation. In contrast, the IO of postnatal day 8 (P8) rats does not exhibit responses to external stimuli. Instead, IO activity primarily reflects corollary discharges associated with the production of self-generated limb twitches during active (REM) sleep. Because corollary discharges are necessary for the computation of internal models, we tested the hypothesis that IO-related corollary discharge is necessary for the emergence of a functioning cerebellar internal model during development. First, by conducting extracellular recordings in the IO at P12 and P20, we confirmed the presence of twitch-related corollary discharge at both ages; however, whereas the IO at P20 responded to limb stimulation, the IO at P12 did not. Next, using a protocol for selectively lesioning the climbing fibers that connect the IO to the cerebellum, including the interpositus nucleus (IP), we confirmed that lesioning at P12 prevents the IP’s expression of corollary discharge at P13. Finally, we assessed the necessity of IO input to the cerebellum for the emergence of an internal model by lesioning climbing fibers at P12 or P19 and testing for the expression of a cerebellar internal model in the thalamus at P20. Only when the lesions occurred at P12 was the expression of the internal model severely disrupted. These findings provide the most direct evidence to date linking twitch-related corollary discharge to the developmental emergence of a cerebellar-dependent internal model.

**SIGNIFICANCE STATEMENT:** In adult animals, internal models of movement generated by the cerebellum are essential for smooth, precise, and adaptive behavior. It is known that these internal models must develop in early infancy in an experience-dependent fashion; however, it is not known how sensorimotor experience acting on the nascent cerebellar circuit gives rise to adult functions. Here, in pre-weanling rats, we show that self-generated signals during sleep provide the early experiences that shape cerebellar internal models. Our findings lay a foundation for understanding how neural activity during sleep shapes typical and atypical brain development.

## INTRODUCTION

Brain development encompasses long periods of continuous change, but periods of discontinuous change also abound (Oppenheim, 1984). For example, the cortical subplate is a transient structure that, before it regresses, scaffolds developing thalamocortical connections (Molnar et al., 2020). In other cases where neural structures do not regress, they exhibit transient functions that are distinct from their canonical functions in adults. For example, primary motor cortex (M1) is an exclusively sensory structure long before it develops the ability to initiate motor commands (Chakrabarty et al., 2000; Dooley & Blumberg 2018; Blumberg & Adolph 2023). Also, the selective but transient suppression of movement-related sensory feedback (i.e., reafference) during wakefulness places an early premium on reafference arising from limb twitches during active (REM) sleep (Tiriac et al., 2016; Dooley & Blumberg 2018). Here, we identify a similar case of provisional development in a precerebellar structure—the inferior olive (IO)—whose canonical functions in awake adult animals depend on external sensory input (Ito et al., 1982; De Zeeuw et al., 1998; Medina & Lisberger, 2008; Herzfeld et al., 2018), but whose preweanling activity in rats reflects a heightened sensitivity to twitch-related reafference during active sleep (Mukherjee et al., 2018). As we show here, this transient feature of IO functioning supports the cerebellum’s ability to build internal models of movement.

In adults, cerebellar internal models enable the computation of sensorimotor predictions. As currently understood, when a self-generated movement is produced, a motor command drives muscle activity; at the same time, a copy of the motor command (i.e., corollary discharge) and reafference from the ensuing movement provide the ingredients for building the internal model (Miall et al., 1993). The output of the internal model is conveyed from the deep cerebellar nuclei (including the interpositus nucleus, IP) to the IO, suppressing expected sensory input at the IO but allowing unexpected input to pass; this unexpected input produces an error signal that is conveyed to the cerebellum via the IO’s climbing fibers to enable updating of the internal model (Devor, 2002; Markov et al., 2021).

In early infancy, it is not possible to distinguish expected from unexpected sensory input because the internal model that enables that distinction has not yet developed. We focus here on the processes that scaffold the developmental emergence of that distinction, particularly the critical role played by corollary discharge. As shown previously in rats at postnatal day 8 (P8), the same motor commands in the midbrain that produce limb twitches during active sleep also produce corollary discharges that are conveyed to the IO (Mukherjee et al., 2018). Importantly, even as the IO conveys corollary discharges to the cerebellum, our initial results suggested that it does not do the same for externally produced sensory stimuli. Because the adult IO is highly responsive to external stimuli (De Zeeuw et al., 1998; Lang et al., 2017; Streng et al., 2018), an early absence of responsivity—if true—requires explanation. That cerebellar-dependent learning begins as early as P17 (Stanton et al., 1992; Campolattaro & Freeman, 2008) and a cerebellar internal model of movement emerges by P20 (Dooley et al., 2021), we hypothesized that a transition in IO functionality occurs toward the end of the third postnatal week.

Here, we test this hypothesis by first recording extracellular IO activity in unanesthetized P12 and P20 rats and document a transition in the IO’s responsivity to externally produced sensory stimuli. Next, by lesioning climbing fibers at P12 and recording IP responses to self- and other-generated movements at P13, we establish the IO as the principal source of corollary discharge to IP at this age. Finally, to test the hypothesis that corollary discharges are necessary for developing cerebellar-dependent internal models (Mukherjee et al., 2018; Dooley et al., 2021), we lesioned climbing fibers at P12 or P19 and, at P20, recorded activity in the ventrolateral thalamus (VL), a primary output of the IP whose twitch-related activity reveals the first functional cerebellar-dependent internal model (Dooley et al., 2021). Altogether, our findings reveal a transient period of development during which the IO preferentially conveys corollary discharges to the cerebellum to scaffold its emerging capacity to compute internal models of movement.

## MATERIALS AND METHODS

All experiments were conducted in accordance with the National Institutes of Health Guide for the Care and Use of Laboratory Animals (NIH Publication No. 80-23) and were approved by the Institutional Animal Care and Use Committee of the University of Iowa.

### Subjects

Male and female Sprague-Dawley rats at P12-13 (hereafter P12; body weight: 32.1 ± 3.2 g) and P19-20 (hereafter P20; body weight: 59.9 ± 7.2 g) were used. Pups were born to dams housed in standard laboratory cages (48×20×26 cm) with a 12 h light/dark cycle. Food and water were available ad libitum. The day of birth was considered P0, and litters were culled to eight pups by P3. Pups were randomly assigned to different age and experimental groups; to protect against litter effects, pups selected from the same litter were never assigned to the same group (Abbey and Howard, 1973; Lazic and Essioux, 2013).

### Surgery

Head-fix surgeries were performed using established methods (Blumberg et al., 2015; Dooley et al., 2021). On the day of testing, a pup with a healthy body weight and a visible milk band was removed from the litter. Under isoflurane anesthesia (3.5–5%, Phoenix Pharmaceuticals), bipolar electrodes (diameter, 0.002 inch; epoxy coated; California Fine Wire) were inserted bilaterally into the nuchal muscle and secured with collodion (Avantor Performance Materials). For pups receiving intramuscular stimulations, an additional bipolar electrode was inserted into the right biceps and secured with collodion. For all pups, the anti-inflammatory agent carprofen (0.1 mg/kg subcutaneous; Putney) was administered. The scalp was sterilized with iodine and isopropyl alcohol, after which the skin on top of the scalp was removed to reveal the skull. The topical analgesic lidocaine hydrochloride (0.2%; Sparhawk Laboratories) was applied to the skull surface and surrounding skin, and Vetbond (3M) was used to secure the remaining skin to the skull. A stainless-steel head-fix (Neurotar) was attached to the skull using a cyanoacrylate adhesive (Loctite, Henkel) that was dried with the accelerant Insta-Set (Bob Smith Industries).

While still under anesthesia, the pup was secured in a stereotaxic apparatus (Stoelting), and a steel trephine (1.8 mm; Fine Science Tools) was used to drill openings in the skull. For IO recordings (P12: n = 8, 4 female; P20: n = 8, 2 female), a hole was drilled on the left side of the skull (i.e., contralateral to the right forelimb; coordinates from lambda, P12: −2.7 mm AP, −0.4 mm ML; P20: −2.5 mm AP, −0.3 mm ML). For IP recordings at P12 (3AP: n = 8, 5 female; saline: n = 8, 4 female), a hole was drilled on the right side of the skull (i.e., ipsilateral to the right forelimb; coordinates from lambda, −1.8 mm AP; +2.0 mm ML). Finally, for VL recordings at P20 (3AP at P12: n = 8, 3 female; saline at P12: n = 8, 4 female; 3AP at P19: n = 8, 3 female) a hole was drilled on the left side of the skull (i.e., contralateral to the right forelimb; coordinates from bregma, −2.1 mm AP, −1.8 mm ML).

Throughout the surgery, the pup’s respiration was monitored, and the concentration of isoflurane was adjusted accordingly. All surgical procedures lasted 16–24 min. After the surgery was complete, the pup was transported to the recording rig, where it was head-fixed in a warm environment between 26.5 and 29°C for at least 1 h before recording began (Dooley et al., 2021). Continuous white noise (70 dB) was present throughout the recording session.

### Data acquisition

To record neural activity at P12, a 16-site linear silicon electrode (10-mm shank; 100 µm between sites; NeuroNexus Technologies) was lowered into the IO (depth at tip: 7.5 ± 0.4 mm). To record neural activity at P20, a 32-site poly silicon electrode (15-mm shank; sites concentrated at the tip; NeuroNexus Technologies) was lowered into the IO. The electrode was coated with fluorescent DiI (Life Technologies) before insertion to enable histological confirmation of the electrode location. Neurophysiological (25 kHz) and EMG (1 kHz) data were acquired using a data acquisition system (Tucker-Davis Technologies). Video was acquired at 100 frames/s using a video camera (Blackfly S, Teledyne FLIR or ace 2 R, Basler). Videos were recorded using SpinView software (Teledyne FLIR) and were synchronized with the electrophysiological data using a custom MATLAB script, as described previously (Dooley et al., 2021).

#### IO activity at P12 and P20

In P12 rats, electrophysiological and video data were recorded for 60 min as the pup cycled freely between sleep and wake. After 60 min, ∼50 intramuscular stimulations of the right forelimb, contralateral to the recording site, were delivered using an isolated pulse stimulator (AM Systems, Model 2100), as described previously (Gómez et al., 2021). Intramuscular stimulations occurred during active sleep or wake and consisted of a single, 20-ms biphasic pulse delivered while the pup was behaviorally quiescent and at least 2 s after the cessation of a limb movement. Pulses were delivered at random intervals of approximately 8 to 20 s. Before recordings began, an initial effective voltage (5.0–9.4 V) sufficient to move the limb was determined for each pup; if, over multiple stimulations, the evoked forelimb movements diminished, the voltage was slowly increased until movements were restored (maximum increase: 0.7 V).

P20 rats were also tested as described above but with one change in the protocol. Because pups exhibit less active sleep at P20 than at P12, the amount of time required to ensure a sufficient number of twitches for analysis is more variable. Thus, we inverted the order of the protocol such that the forelimb stimulations occurred first, followed by 3-4 h of sleep-wake recording.

#### IP activity at P13 after climbing-fiber lesions

Using a method described previously (Jones et al., 1994), climbing fibers were lesioned at P11-12 using successive injections of harmaline (Thermo Scientific), 3 acetylpyridine (3AP, Sigma-Aldrich), and nicotinamide (Sigma-Aldrich). Harmaline was dissolved in DMSO and diluted to 10% in saline, and 3AP was diluted to 9.8% in saline. Over the 3.5-h injection protocol, pups were housed in an incubator at an air temperature of ∼37°C. Pups were first injected intraperitoneally with harmaline (2 µl/g body weight). Approximately 1 h later, when body tremors began, the pups were injected intraperitoneally with 3AP (6.8 µl/g body weight). The tremors produced by harmaline reflect excitation of climbing fibers, and the 3AP targets those fibers for lesioning (Anderson et al., 1987). Approximately 2 h after injection of 3AP when the harmaline-induced tremors subsided, nicotinamide was injected intraperitoneally (7.5 µl/g body weight) to inhibit further action of 3AP (Jolicoeur et al., 1982). Control animals were given equivalent volumes of saline at all three injection times. Pups were then returned to their litters until testing, which always occurred 1 day later. At that time, using the methods described above, IP activity was recorded across sleep-wake cycles and during forelimb stimulations.

Histological assessment of the effectiveness of the pharmacological lesions was conducted in four additional P12 rats. Two pups experienced the lesion protocol and two the control procedure. Then, on P13, each pup was anesthetized with 2–5% isoflurane and secured in a stereotaxic apparatus as described above. A hole was drilled on the right side of the skull over the cerebellum (coordinates from lambda: −1.8 mm AP; +2.0 mm ML), and a 0.5 µL syringe (Hamilton) containing 0.05 µL of 2% wheat germ agglutinin (WGA) conjugated to Alexa Fluor 555 (WGA-555; Invitrogen Life Technologies) was inserted into the cerebellar cortex (DV: −2.0 mm). After 10 min, WGA was slowly injected (∼0.1 µL per min). Then, after another 10 min, the microsyringe was withdrawn and the incision was closed with Vetbond (3M). The pup was returned to its home cage and perfused 24 h later as described below.

#### VL thalamus activity at P20 after climbing-fiber lesions at P12 or P19

The protocol for producing climbing-fiber lesions and controls was performed at P12 or P19 as described above. Then, at P20—that is, eight days or one day later, respectively—VL activity was recorded using the method described above across sleep-wake cycles and during forelimb stimulations.

### Histology

At the end of the recording sessions, pups were anesthetized using an intraperitoneal injection of ketamine-xylazine (90:10; 0.1 ml/kg b.w; Akron). Pups were perfused transcardially with phosphate-buffered saline followed by 4% paraformaldehyde (PFA). Brains were extracted and placed in PFA for at least 24 h, after which they were transferred to phosphate-buffered sucrose for at least 48 h before coronal sectioning (80 µm). A fluorescent microscope (Leica Microsystems) was used to visualize electrode locations before staining with Cresyl violet.

### Data analysis

#### Spike sorting

Electrophysiological and behavioral data were imported into MATLAB and analyzed as described previously (Gómez et al., 2023). Custom scripts were used to filter raw neurophysiological data (bandpass: 500-5,000 Hz) and individual units were extracted using Kilosort (Pachitariu et al., 2016) and Phy2 (Rossant and Harris, 2022).

##### Video analysis of movement and behavioral-state determination

Electrophysiological data and video data were imported into Spike2 (Cambridge Electronic Design). Behavioral states were identified based on nuchal EMG activity and behavior from video (Del Rio-Bermudez et al., 2016; Gómez et al., 2023). For this analysis, we focused on periods of active wake and active sleep. Active wake was defined as a period of high muscle tone during which the pup exhibited coordinated and/or high-amplitude limb movements. Active sleep was defined as a period of muscle atonia punctuated by sharp spikes in the nuchal EMG record indicative of limb twitches. The remaining periods, consisting of behavioral quiescence that could indicate either quiet wake or quiet sleep, were not analyzed.

Limb movements were detected using a custom MATLAB script that identified frame-by-frame changes in pixel intensity within regions of interest (ROIs; Dooley et al., 2020). Using an ROI encompassing the right forelimb, the number of pixels with >5% changes in pixel intensity were summed frame by frame, resulting in a trace of real-time movement. Using a threshold for displacement of the right forelimb, events were triggered at movement onset and subsequently confirmed visually by the experimenter from video. For IO and IP recordings, mean onset of limb movements was calculated over all triggered twitches and stimulations (Richardson et a., 2024); for VL thalamus recordings, median displacement of the limb was calculated and normalized over all triggered twitches (Dooley et al., 2021). Finally, the onset of each intramuscular limb stimulation was designated as the video frame immediately preceding the evoked limb movement. Stimulations that evoked global increases in movement were excluded from analysis.

##### Determination of responsive and non-responsive units

Units were classified as either responsive or nonresponsive to a twitch or intramuscular stimulation (Richardson et al., 2024). Briefly, baseline activity was established in the period prior to stimulus onset, and individual units were considered responsive if the mean post-stimulus peak firing rate was more than 4 standard deviations above baseline.

##### Width at half-height

For responsive units only, the width at half-height for each response profile was determined. After normalizing each unit’s peak response in relation to baseline activity, a five-bin kernel was used to smooth the data before interpolating the data to increase temporal precision (from 10-ms to 1-ms bins using the MATLAB interp1() function). The width at half-height was defined as the duration of the response profile (in ms) at the halfway point between baseline and peak amplitude.

### Statistical analysis

The Shapiro–Wilk test for normality and the Levene’s test for homogeneity were conducted before analysis. When either of these tests failed, Welch’s t test was used instead. When proportions were tested, an arcsine transformation was used. Between-subject t tests were performed to compare the mean widths at half-height, peak response times, and change in amplitude for stimulus-responsive units. All statistical tests were performed using SPSS (IBM). Unless otherwise noted, means are presented with standard errors, and alpha was set at 0.05. After ANOVAs were calculated, and when appropriate, a Tukey’s (in the case of equal variances) or Games-Howell (in the case of unequal variances) post hoc test was conducted. When appropriate, a Bonferroni correction was used to correct alpha for multiple comparisons. Effect size was calculated using an adjusted partial eta-squared for ANOVAs (adj ηp 2) and adjusted eta-squared for t tests (adj η2; Mordkoff, 2019).

## RESULTS

We recorded extracellular IO unit activity at P12 and P20 in head-fixed rats (Fig. 1*A*). Histology confirmed electrode placement in the IO (Fig. 1*B*). Pups cycled freely between sleep and wake throughout the recording sessions. Representative recordings of IO unit activity and nuchal EMG in relation to right-forelimb twitches during active sleep and right-forelimb stimulations are shown in Figure 1*C-D*. Also shown are IO units that were classified as responsive or nonresponsive to twitches or stimulations (see Materials and Methods, Determination of responsive and nonresponsive units).

**Figure 1.**
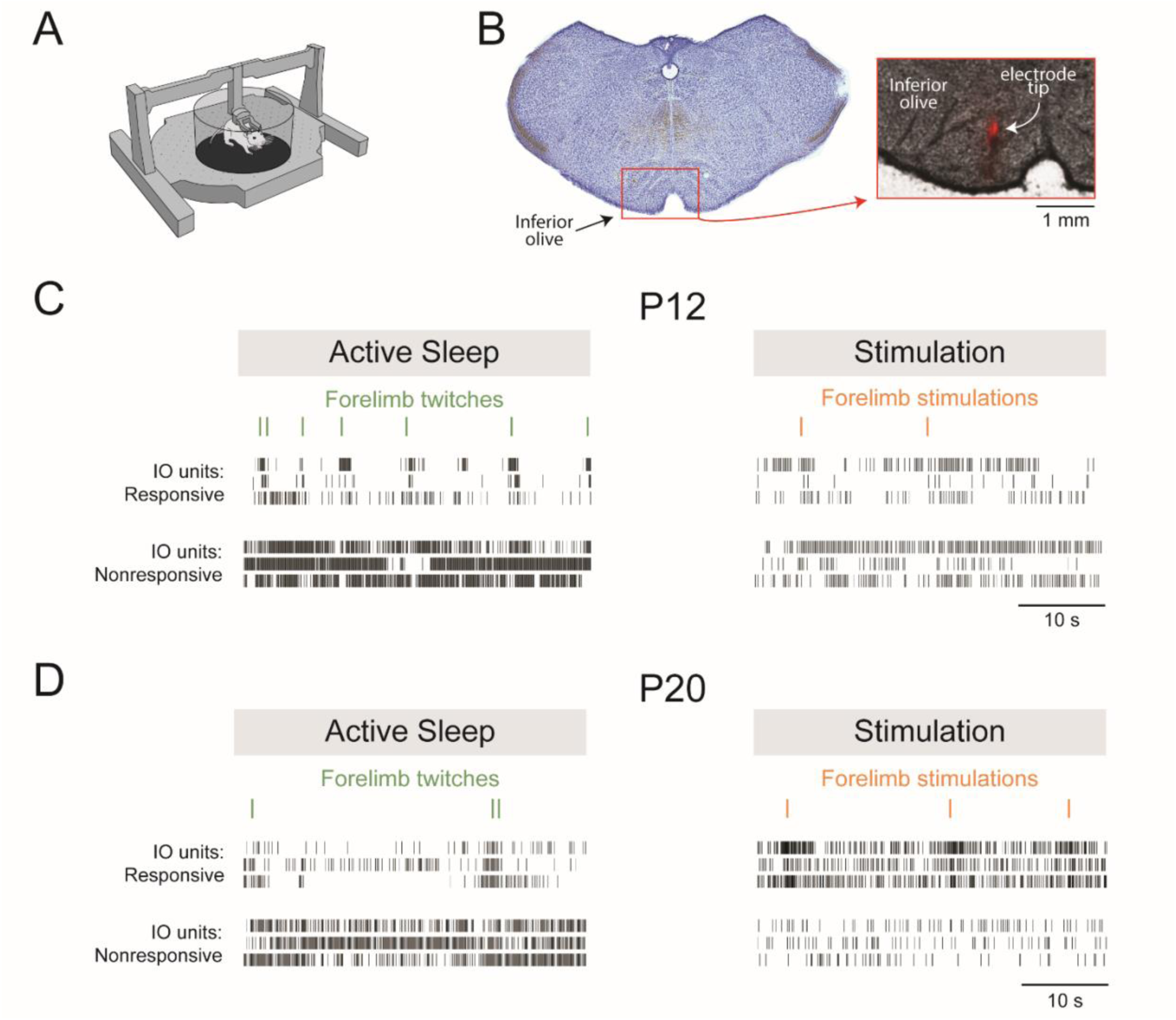
Experimental design and representative data. ***A***, Illustration showing a head-fixed pup in the recording apparatus. ***B***, Histology showing the red fluorescent electrode track in the IO. ***C***, Representative data from P12 rats showing IO responses to forelimb twitches (left) and responses to intramuscular electrical stimulation of the forelimb (right). From top: ticks denoting the onset of a forelimb movement (twitches in green; stimulations in orange, and IO activity for units that were responsive or nonresponsive to twitches or muscle stimulations. IO recordings were contralateral to the right forelimb. ***D***, Same as in ***C*** except for P20 rats.

### Developmental emergence of exafferent activation of the inferior olive

The twitch-related response profiles in Figure 2*A* exhibit a sharp peak at P12 compared with the broader peak at P20, suggesting a change in the nature of the input to the IO at these two ages. Overall, the proportion of all IO units that were responsive to twitches was higher at P12 (59%, n=88 units) than at P20 (40% n=95 units). However, when the proportions were averaged within each pup, the difference was not significant (*t*_(14)_=2.03; Fig. 2*B*, *left*). There was also no significant difference in the peak firing rate of responsive units per pup across ages (*t*_(14)_=0.44; Fig. 2*B*, *right*).

**Figure 2.**
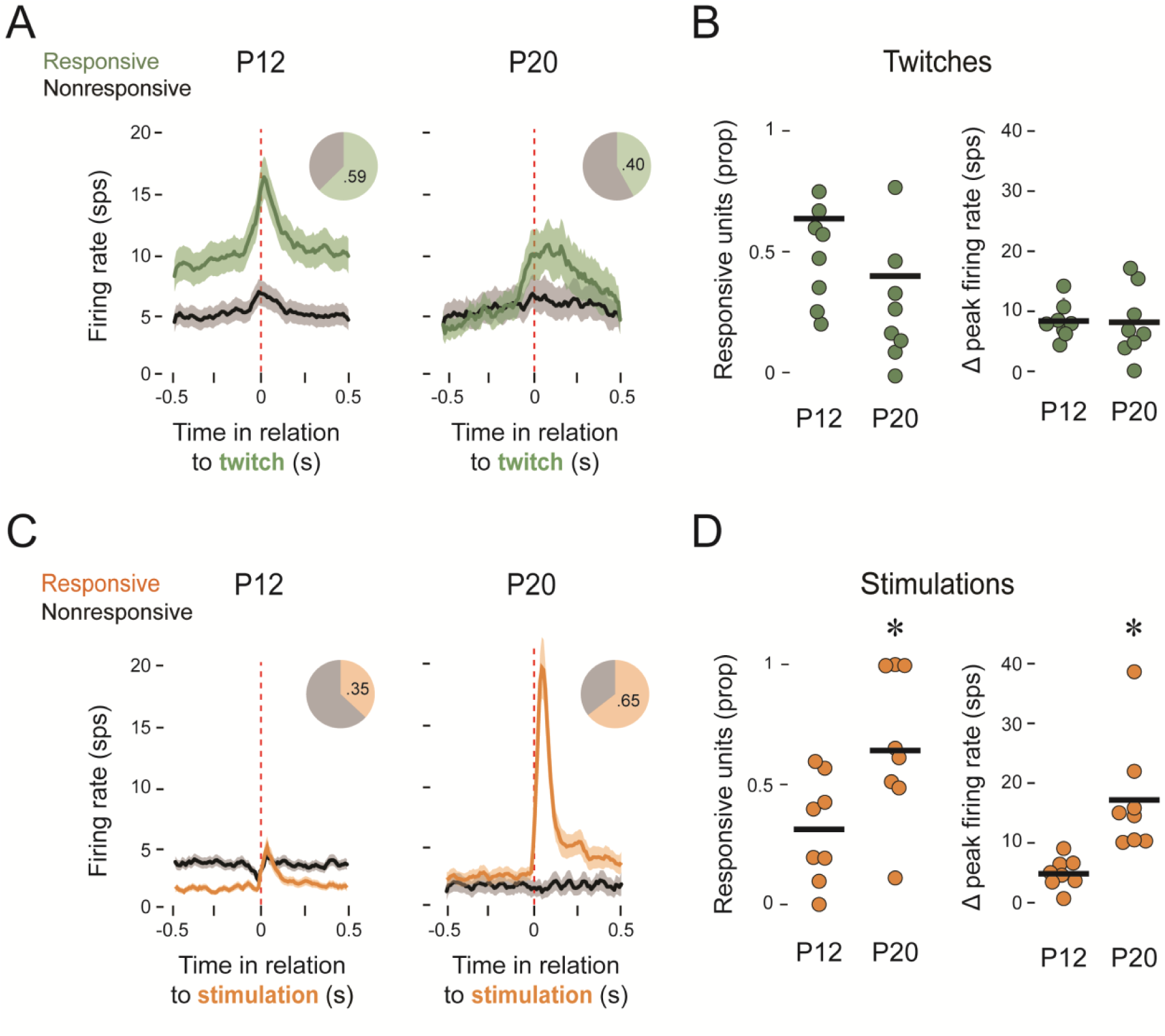
IO responses to twitches and stimulations at P12 and P20. ***A***, Perievent histogram of IO activity for responsive (green) and nonresponsive (black) units in relation to the onset of forelimb twitches at P12 (left; n=8 pups and 86 total units) and P20 (right; n=8 pups and 95 total units). Pie charts show the proportion of units that were responsive. ***B***, Left: Individual and group mean proportions of twitch-responsive IO units at P12 and P20. Right: Individual and group mean changes in peak firing rate from baseline for each pup for units responsive to twitches at P12 and P20. ***C***, Same as in ***A*** but for unit activity in response to forelimb stimulations (orange). ***D***, Same as in ***B*** but for stimulation-responsive units. The asterisks denote significant differences between groups.

For stimulations, the large, sharp peak in the IO response at P20 dwarfs the response at P12 (Fig. 2*D*); also, a significantly higher proportion of units were responsive at P20 (65%) than at P12 (35%). When average proportions were calculated for each pup, this difference was significant (*t*_(14)_=2.7; p=0.017; adj ⴄ^2^=1.408; Fig. 2*E*, *left*). Similarly, there was a significant increase in the peak firing rate of responsive units per pup between P12 and P20 (Welch’s *t*_(7.90)_=3.65; p=0.006; adj ⴄ^2^=1.32; Fig. 2*E*, *right*). These results clearly demonstrate the emergence of IO responsivity to external stimulation over the third postnatal week.

### Developmental emergence of twitch-related reafference in the inferior olive

To further quantify the age-related change in the twitch-related response profiles in Figure 2*A*, we measured the width at half-height and the timing of the response peaks for each individual unit. The response profiles of the 49 responsive units at P12 were uniform among twitch-responsive units and the timing of the response peaks occurred within 50 ms of the onset of twitches, indicative of corollary discharges (Fig. 3*A*; Mukherjee et al., 2018). By P20, the response profiles for the 39 responsive units were more variable; at this age, although the response profiles of some units continued to peak near the onset of twitches, more units were peaking after 100 ms of twitch onset, indicative of reafference (Fig. 3*B*). Whereas the widths at half-height, averaged within each pup, were not significantly different between P12 and P20 (*t*_(14)_= 1.54; Fig. 3*C*), the times of the response peaks were significantly longer at P20 (*t*_(14)_=2.55; p=0.023; adj ⴄ^2^=0.310; Fig. 3*D*). Thus, in addition to the increased responsivity of the IO to exafferent stimulation between P12 and P20, its twitch-related activity becomes more complex—reflecting input from both reafference and corollary discharge.

**Figure 3.**
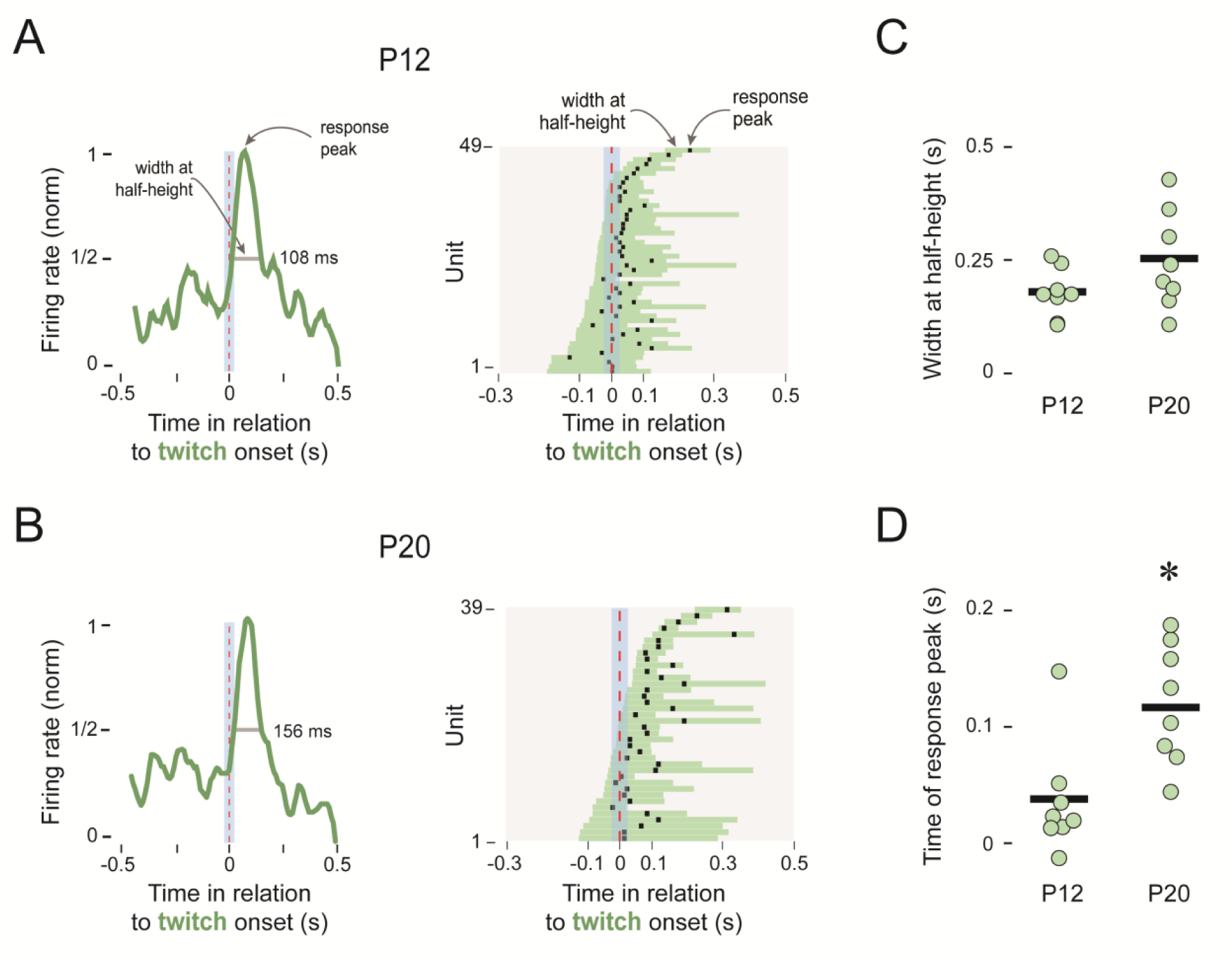
Twitch-related response profiles for the IO at P12 and P20. ***A***, Left: Representative perievent histogram (normalized) for a twitch-responsive unit at P12. Width at half-height and time of peak response are indicated. Right: Widths at half-height (horizontal bars) and times of response peak (black squares) for all 49 twitch-responsive units at P12. ***B,*** Same as in ***A*** but for all 39 twitch-responsive units at P20. ***C,*** Individual and group mean widths at half-height for twitch-responsive units at P12 and P20. ***D,*** Same as in ***C*** but for time of response peak. The asterisk denotes a significant difference between groups.

### Climbing-fiber lesions at P12 disrupt the conveyance of corollary discharge from inferior olive to interpositus

Recently, we demonstrated in P12 rats that the IP responds robustly to twitch-related reafference but not exafference (Richardson et al., 2024). We hypothesized that the IO is a critical factor in the activity profile of IP, perhaps due to the exuberant climbing-fiber collaterals at that age (Raman & Najac, 2017; Pickford & Apps, 2017). To test this hypothesis, we produced pharmacological lesions that target climbing fibers (Fig. 4*A*) and recorded from the IP one day later (Fig. 4*B*). We confirmed the efficacy of the lesion method at P12 by injecting a fluorescent retrograde tracer (WGA) into the cerebellar cortex (Fig. 4*C*). Because the cerebellar cortex receives abundant climbing-fiber projections, effective lesions would prevent the tracer from traveling to cell bodies in the IO. To test the specificity of our lesion, we also assessed the presence of fluorescent cell bodies in the lateral reticular nucleus (LRN), which also projects to cerebellar cortex and conveys twitch-related activity to the cerebellum (Mukherjee et al., 2018;Font et al., 1997). In the control pup, the WGA tracer labeled cell bodies in both the IO and the LRN (Fig. 4*C*). In contrast, the tracer in the lesioned pup only labeled cell bodies in the LRN; the IO was devoid of labeled cell bodies. Thus, our lesion method selectively lesioned climbing fibers from the IO.

**Figure 4.**
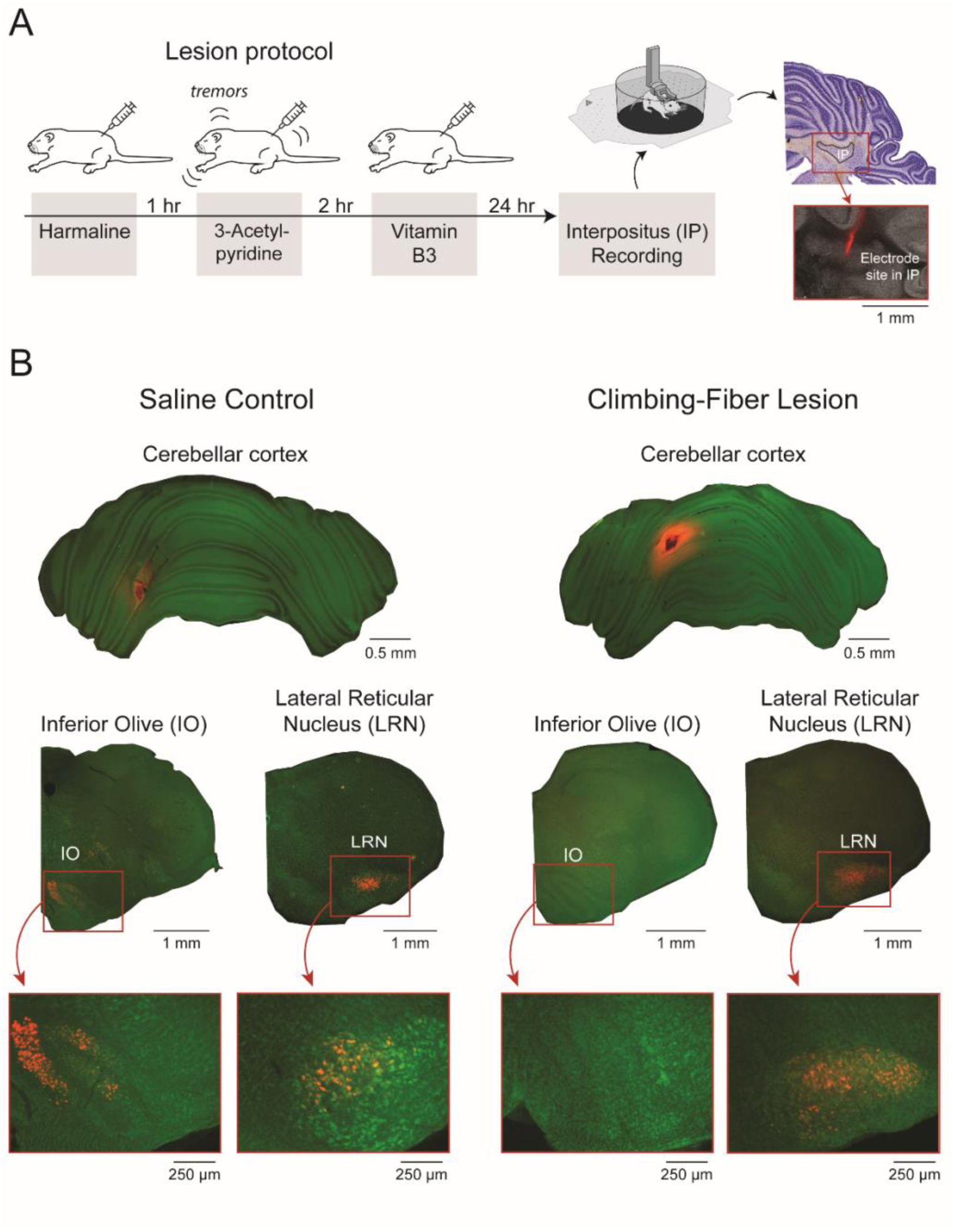
Protocol for producing climbing-fiber lesions and recording from IP. ***A,*** Timeline of lesion protocol and histology showing the red fluorescent electrode track in IP. ***B,*** WGA expression (red) for control (left) and climbing-fiber lesioned (right) pups. Top row: WGA injection site (red) in cerebellar cortex. Middle row: WGA expression in inferior olive (IO) and lateral reticular nucleus (LRN). Bottom row: 5x magnifications of WGA expression for the sections above.

In the lesion and control pups at P13, 71-73% of the IP units were twitch-responsive (Fig. 5*A*). Neither the proportion of responsive units per pup (*t*_(14)_= 1.74; Fig. 5*B*) nor the width at half-height (*t*_(14)_= 1.41; data not shown) differed significantly between the two groups. In contrast, there was a significant difference in the peak response time (*t*_(14)_= −2.63, *p* = 0.019, adj ⴄ^2^=1.516), with the average peak time shifting to the right for the lesion group (Fig. 5*D*). This delayed peak in the lesioned pups indicates a shift in IP responding away from corollary discharge and toward sensory responding when no longer receiving input from IO.

**Figure 5.**
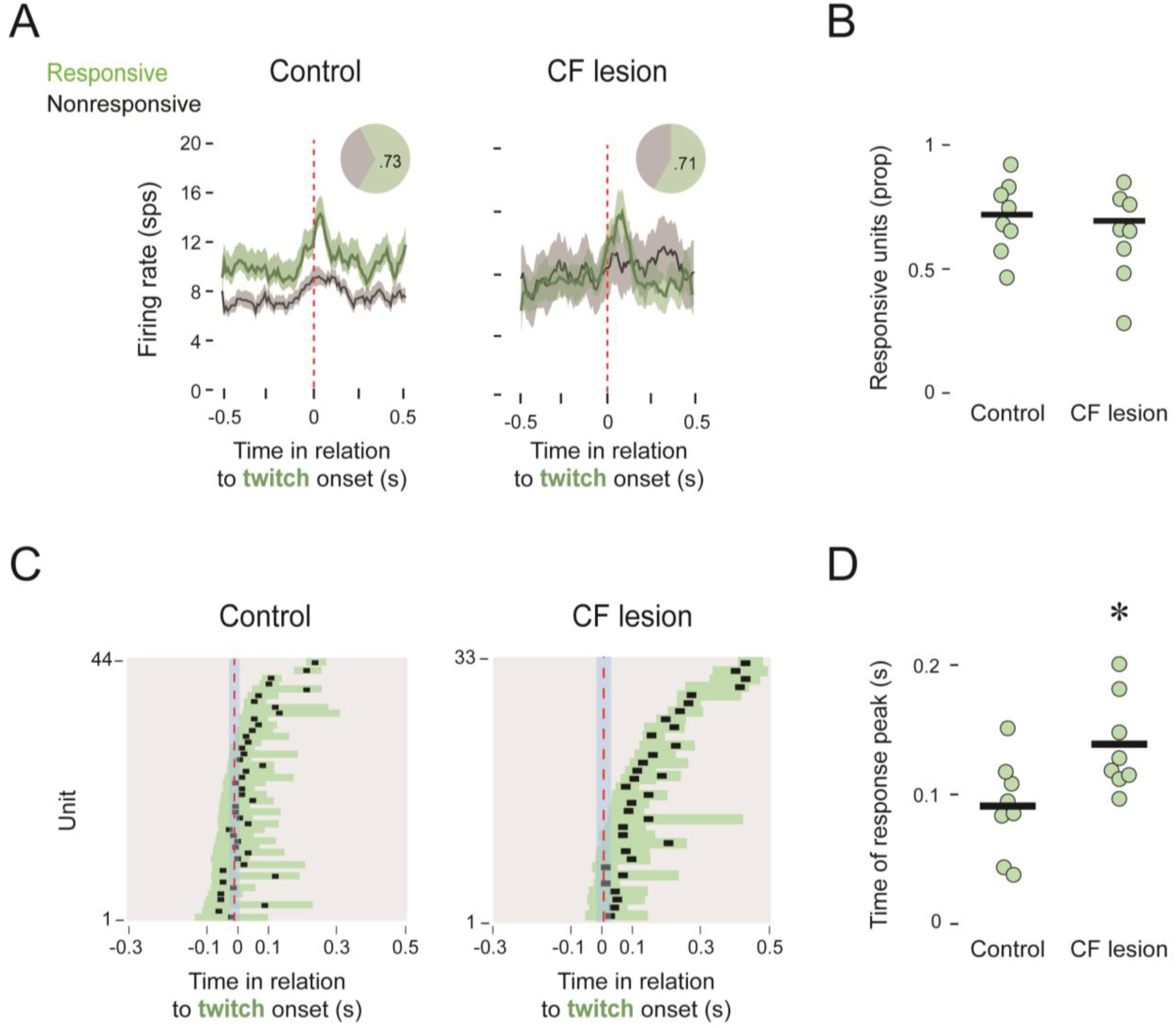
Twitch-related IP activity after climbing-fiber (CF) lesions. ***A,*** Mean (±SEM) perievent histograms of IP unit activity for responsive (green) and nonresponsive (black) units in relation to the onset of forelimb twitches control (left; n=8 pups, 60 units) and climbing-fiber lesions (right; n=8 pups, 46 units). Pie charts show the proportion of responsive units. ***B,*** Individual and group mean proportions of twitch-responsive IP units in control and lesion groups. ***C,*** Widths at half-height (horizontal bars) and times of response peak (black squares) for all individual twitch-responsive units for control (left) and lesioned (right) pups. ***D,*** Mean time of response peak for each pup in the control and lesion groups. The asterisk denotes a significant difference between groups.

### Climbing-fiber lesions at P12 disrupt the emergence of the internal model

We next tested the hypothesis that the corollary-discharge signal from the IO is necessary for the emergence of a cerebellar-dependent internal model between P12 and P20 (Dooley et al., 2021). In P20 rats with and without climbing-fiber lesions, we recorded from VL (Fig. 6*A*) because the degree to which it matches the movement profile of forelimb twitches (Fig. 6*B*) provides an assay of the precision of the internal model. In addition to the lesion and control pups at P12, we added a group of lesioned pups at P19 to assess the acute effects of climbing-fiber lesions on expression of the internal model at P20.

**Figure 6.**
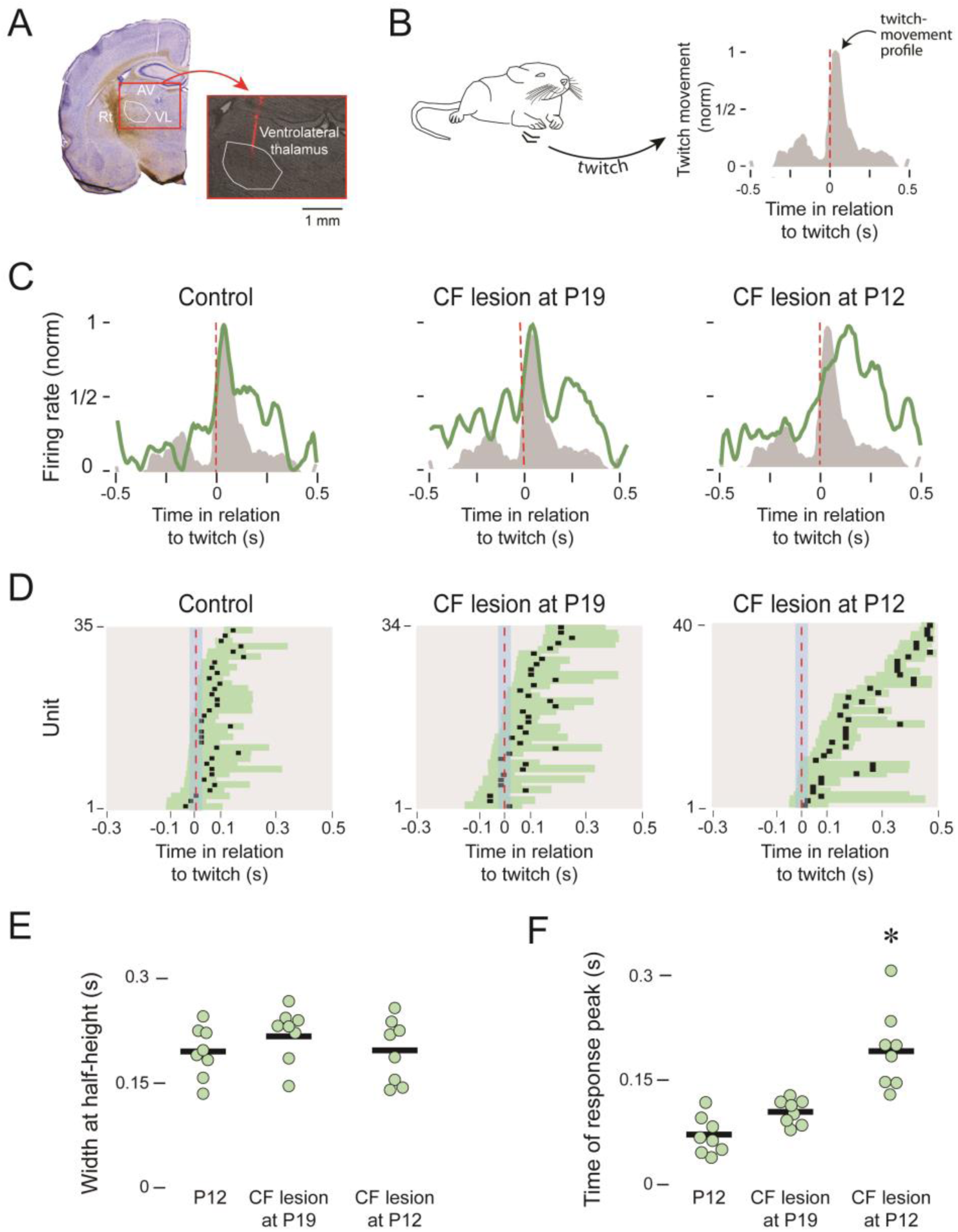
Twitch-related VL activity after climbing fiber (CF) lesions. ***A,*** Histology showing the red-fluorescent track of the recording electrode in VL. Adjacent thalamic regions are also labelled. Rt: thalamic reticular nucleus; AV: anteroventral nucleus. ***B,*** Average movement profiles (normalized) for the right forelimb in relation to twitch onset. ***C,*** Representative perievent histograms (normalized) showing VL activity (green) in relation to twitch-movement profiles (grey) for control pups (left) and pups receiving CF lesions at P19 (middle) and P12 (right). ***D,*** Widths at half-height (horizontal bars) and times of response peak (black squares) for all forelimb-responsive units for control pups (left) and pups receiving CF lesions at P19 (middle) and P12 (right). ***E,*** Individual and group mean times of response peak for twitch-responsive units. ***F,*** Same as in E but for widths at half-height. The asterisk denotes a significant difference from the other two groups.

Representative response profiles of VL units in each of the three groups illustrate how VL activity matches the movement profile of twitches in a control pup and in a pup with a climbing-fiber lesion at P19, but in the pup with climbing fiber lesions at P12 the VL peak is substantially delayed (Fig. 6*C*). These representative profiles are consistent with the peak response times across all units in the three groups (Fig. 6*D*), in which the mismatch between the twitch and the neural response are only evident in the pups with P12 lesions. Although there was no significant main effect of group on width at half-height (*F*_(2,21)_ = 1.13; Fig. 6*E*), there was a significant main effect of group on peak response time (*F*_(2,21)_ =10.97, *p* = 0.001, adj *η_p_^2^* = 0.577; Fig. 6*F*). Post-hoc tests revealed a significant difference between the P12 lesion group and the other two groups (controls: *p* = 0.0034; P19 lesions: *p* = 0.0154). Thus, these results demonstrate the necessity of IO input to the cerebellum for the developmental emergence of the internal model between P12 and P20, but that the acute loss of IO input only one day earlier exerts a minimal effect on the model’s expression.

## DISCUSSION

With the onset of weaning around the third postnatal week, rats increasingly gain independence from their mother as their behavioral repertoire expands (Cramer et al., 1990; Thiels et al., 1990). It is around this time, at P20, that we see the emergence of a cerebellar internal model for movement (Dooley et al., 2021). Also, as shown here, P20 marks the end of the IO’s sensory isolation: Whereas the IO at P12 primarily responds to the corollary discharge component of twitches, by P20 the IO’s response profile has expanded to include reafference and exafference (Fig. 7). What is the functional significance of the IO’s sensory isolation and associated accentuation of corollary discharge? We answered this question by showing that climbing-fiber lesions at P12 prevent the typical development of the cerebellar internal model by P20. Together, these results identify a sensitive period when IO activity sculpts the cerebellar network’s ability to make accurate predictions about self-generated movement.

**Figure 7.**
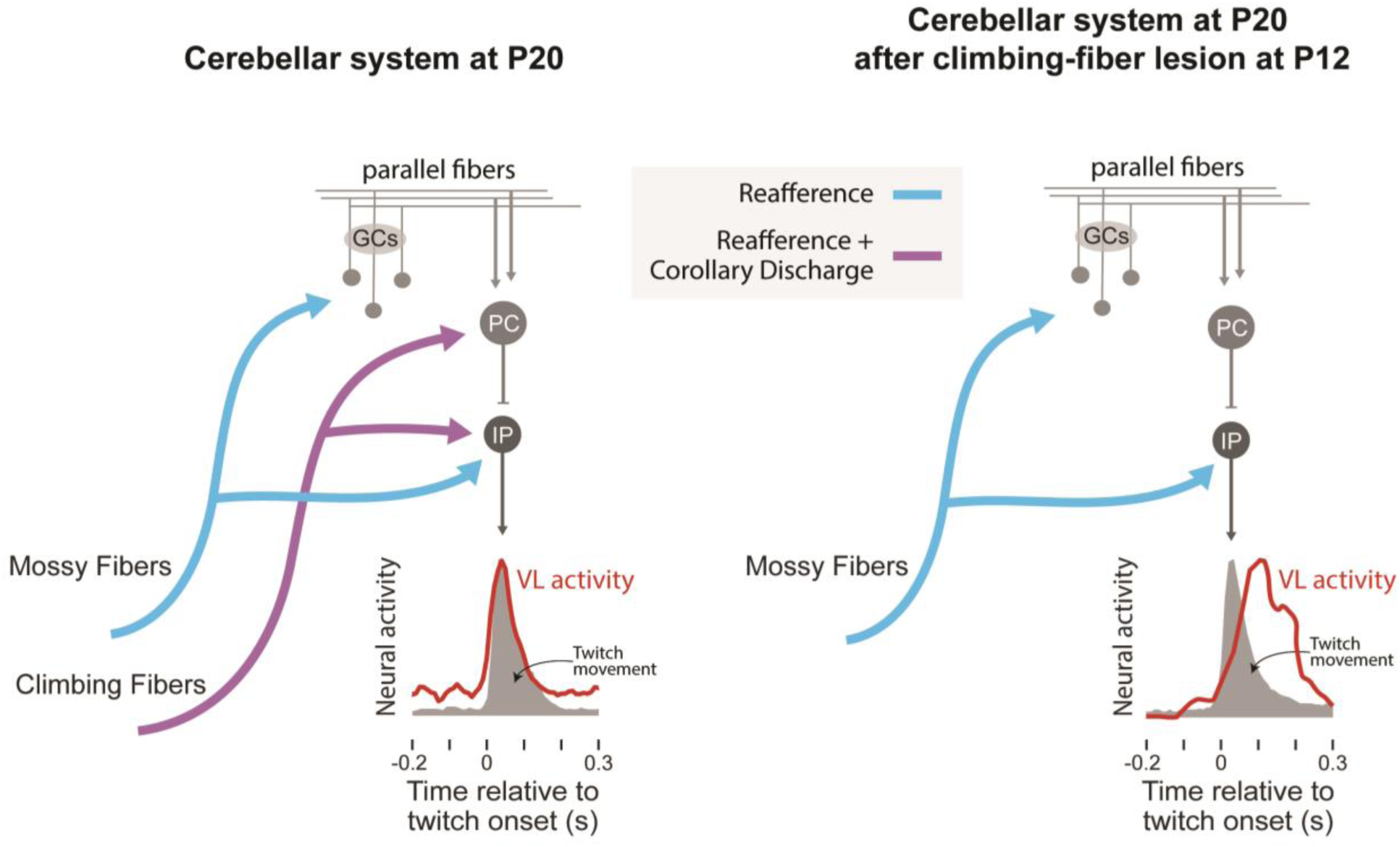
The developmental emergence of a cerebellar internal model is dependent on corollary discharge from the IO. Left: In typically developing rats, climbing fibers arising from the IO and mossy fibers arising from other precerebellar nuclei convey twitch-related corollary discharge (purple lines) and reafference (blue lines), respectively. The emergence of the cerebellar internal model between P12 and P20, as expressed by twitch-related activity in ventrolateral thalamus (VL), relies on the convergence of both kinds of signals in the cerebellum. Right: After lesions of climbing fibers at P12 and the resulting loss of corollary discharge over the ensuring week, the internal model does not develop. Corollary discharge signals are also conveyed to the cerebellum via mossy fibers from the lateral reticular nucleus; those inputs are not shown (see Discussion). GCs: granule cells; PC: Purkinje cell; IP: interpositus.

### Transient sensory gating shapes cerebellar development

When Mukherjee and colleagues (2018) recorded self- and other-generated activity in precerebellar nuclei, they saw that IO activity in infants did not function like that in adults, that is, as a sensory-driven error detector (Ito, 2008; Raymond & Medina, 2018). Specifically, at P8, it seemed that sensory responses could not be elicited from the IO. However, this observation was only preliminary. Thus, here we systematically investigated this phenomenon at P12 and found that this lack of sensory responsiveness was indeed reliable. But why would the cerebellar system exhibit such profound sensory isolation at these early ages? We propose that this isolation is a developmental strategy in that it creates conditions necessary for building internal models.

To build internal models, the cerebellum must calculate the delay between the arrival of a corollary discharge and its corresponding reafferent signal. This calculation must be performed for each limb because feedback delays differ depending on physical distance; for example, forelimb signals arrive at the cerebellum more quickly than hindlimb signals. Additionally, feedback delays change as pups triple in size over the first three postnatal weeks. Therefore, the cerebellar system must precisely and flexibly distinguish between corollary discharges and reafference. Routing corollary discharge through climbing fibers provides a salient temporal marker—a complex spike—that can be compared with later-arriving mossy-fiber input (Catz et al., 2005). When we lesioned climbing fibers at P12, that comparison could not be made and internal models failed to form. It should be noted that although the LRN also conveys corollary discharge (Mukherjee et al., 2018), and although its projections to the cerebellum were spared by our lesion method, those projections seem to be insufficient for development of the internal model. Thus, combining the current findings with those from our earlier report (Richardson et al., 2024), we conclude that at these early ages corollary discharge arising from the IO is what drives development of the cerebellar-dependent internal model.

Through what mechanism is the IO at P12 isolated from peripheral sensory stimuli? Although it is known that inhibitory feedback from the cerebellar cortex silences exafference in the interpositus nucleus (Richardson et al., 2024), it cannot explain the lack of sensory responsiveness in the IO. One possibility is that subthreshold oscillations among electrically coupled IO neurons help to gate sensory input (Giessen et al., 2008; Stern et al., 2011; Loyola et al., 2023), but similar gating in other precerebellar nuclei (Freeman & Muckler, 2003; Campolattaro & Freeman, 2008) suggests a more general mechanism that remains unknown.

### Implications for the maintenance of internal models into adulthood

Even though the IO at P20 has begun to respond to peripheral stimuli, it continues to process twitch-related corollary discharge. This observation raises the question of whether this responsivity to corollary discharge continues into adulthood and, if it does, if it continues to exert the same influence on IO activity. However, in most adult models of cerebellar function, corollary discharge arises during wakefulness from motor cortex via the pons. In turn, the cerebellum receives the corollary discharge from the pons and creates a prediction signal that is conveyed to the IO where it suppresses expected input (Ito, 1984; Devor, 2002; Medina et al., 2002). Thus, there remain significant gaps in understanding how our observations of state-dependent IO activity connect with what is known in adults during wakefulness.

Cerebellar internal models are primarily updated by ‘error-correction’ signals from the IO (Medina & Lisberger, 2008). These signals have only been studied in the context of wake behaviors. However, if twitching continues to produce corollary discharge in the adult IO, then it is possible that twitch-related IO activity contributes to cerebellar learning processes, including the integration of cerebellar internal models with forebrain circuits (Walker, 2006).

After climbing-fiber lesions at P19, we observed little effect on the functioning of the internal model at P20. These control lesions were therefore effective for showing that the internal model does not depend on real-time input once it has developed. However, this finding should not be interpreted to mean that the internal model is impervious to such lesions beyond P20. For example, in the visuomotor system—which also relies on an internal model—surgical disruption of sensory feedback in monkeys and humans causes a slow decay of model accuracy over a period of weeks (Steinback 1986; Lewis et al., 1994; see Wang et al., 2007). Accordingly, in rats after climbing-fiber lesions at P19, we would expect to see a slow decay of model function that would evolve over days or weeks, consistent with long-term plastic changes at the neural level. Whether the development of the internal model depends on the same processes that maintain the internal model at later ages has yet to be resolved.

### Limitations

Twitches are a meaningful class of self-generated movement because they are abundant in early life and, because they are discrete, can readily be associated with neural activity. In contrast, wake movements are noisy and thus more difficult to associate with neural activity. In addition, movements during wake are transmitted to the caudal brainstem and cerebellum, resulting in substantial artifact. Thus, our findings here are necessarily restricted to twitch-related activity and we can only speculate about the possible contributions of wake-related activity.

### Conclusion and future directions

Previously, we discovered that IP fails to respond to externally generated signals at P12 (Richardson et al., 2024). We now find that the same is true for the IO, which is upstream of IP at this age (Nicholson & Freeman, 2003). This suppression of external signals—coupled with robust corollary discharge and reafference—reveals a developmental bias in favor of processing signals associated with self-generated movement. Whether this preference extends to waking behavior or is confined to the self-generated twitches of active sleep remains an open question.

What is clear, however, is that the current findings fit within a general theme characterized by transient state-dependent gating of sensory input in the cerebellar system (Campolattaro & Freeman 2005; Tiriac et al., 2014; Tiriac & Blumberg, 2016; Mukherjee et al., 2018; Dooley & Blumberg, 2018; Richardson et al., 2024). Before the animal can make sense of the sensory barrage of the outside world, it must first learn the sensory consequences of its own movements. Through this self-generated activity, the cerebellum constructs internal models that link motor commands with expected feedback (Ito & Kano, 1982; Jörntell & Hansel, 2006). Thus, there is a rough correspondence between the onset of the cerebellar internal model and increasing responsivity of the cerebellar system to external stimuli. Whether these two complementary processes are causally related is not yet known.

A focus on adult function can blind us to critical but often transient developmental features. Here we have shown that the infant IO exhibits transient features that play a critical role in enabling the cerebellar system to achieve adult functionality. Revealing these features required that we focus on twitches during REM sleep. Thus, studying cerebellar activity in infants—when REM sleep is the predominant behavioral state—reveals unique processes that drive the development of internal models and that could, we propose, also be important for the maintenance of internal models in adults. Consideration of this proposal will lead to a cohesive understanding of sleep’s role in promoting cerebellar plasticity across the lifespan.

## Acknowledgments

This research was supported by a grant from the National Institutes of Health (R37-HD081168) to M.S.B.

